# Parallel lipid-carried epitope pillars enhance stimulation of chimeric antigen receptor T cells

**DOI:** 10.1101/2022.04.15.488352

**Authors:** Jiao Tong, Hanyu Lai, Jingxia Wang, Chenxi Zhu, Xinjia Mai, Dapeng Zhou

**Affiliations:** Tongji University School of Medicine, Shanghai, 200092, China

**Keywords:** Chimeric antigen receptor T cells, glycosylphosphatidylinositol-anchored protein, Glycosylation, Tumor antigen

## Abstract

Chimeric antigen receptor T cells are genetically engineered to express a specific T cell receptor of interest, such as to target a cancer-specific antigen. Signaling events in chimeric antigen receptor T cells are essential for their proliferation, survival, and function. To achieve optimal antitumor efficacy, signaling motifs as well as the structure of the transmembrane domain of the chimeric antigen receptor have to be carefully designed. However, it remains unclear whether the arrangement, and therefore the movement and elasticity, of tumor antigens influence their stimulation of chimeric antigen receptor T cells. Here we show that MUC1 molecules tethered to a glycosylphosphatidylinositol anchor drastically increased its stimulation of chimeric antigen receptor T cells. In particular, MUC1 molecules containing one tandem repeat sequence showed significantly greater stimulatory activity than five tandem repeat sequences. Thus glycosylphosphatidylinositol-anchored, tumor antigen epitope pillars in parallel are attractive targets for the design of chimeric antigen receptor T cells. These novel findings, which we propose as the parallel lipid-carried epitope model, emphasize the importance of epitope arrangement in selecting highly effective chimeric antigen receptor T cells, potentially revolutionizing the applicability of this therapy. Furthermore, our data have implications for the necessity of methodologies to measure the elasticity, movement, and density of antigen pillars in parallel, as key tools to guide future chimeric antigen receptor T cell therapy.

## Introduction

Chimeric antigen receptor T cells are genetically engineered to express a specific T cell receptor of interest, such as to target a cancer-specific antigen. The signaling events for chimeric antigen receptor T cells determine their proliferation, function, and survival.^1^ Chimeric antigen receptor T cells use signaling pathways common to conventional T cells but also have unique signaling events. The T cell receptor/CD3 complex and chimeric antigen receptors share the CD3ξ intracellular region but differ in extracellular antigen-binding domains. A single chimeric antigen receptor molecule may function as a T cell receptor signaling complex containing T cell receptor αβ, CD3ξξ, CD3εδ, and CD3εγ. The mechanisms of T cell receptor activation may include receptor clustering, mechanosensing, and protein segregation that lead to changes in local kinase/phosphatase balance. The chimeric antigen receptors and T cell receptor use similar molecular machinery to transduce signaling, including PLC, Zap-70, LAT, and Src family kinases.^2-3^ However, T cell receptor signaling is much more sensitive than the chimeric antigen receptor because hundreds of antigens are required to activate chimeric antigen receptors.^4^

Activation of both the T cell receptor and chimeric antigen receptor requires the formation of stable contact, the “immunological synapse.” Immunological synapse formation in chimeric antigen receptor T cells is still poorly understood, although it is generally accepted that the quality of immunological synapse formation is essential for chimeric antigen receptor T cell function. The immunological synapse pattern formed by chimeric antigen receptor T cells is significantly different from that formed by T cell receptor T cells. The immunological synapse of the chimeric antigen receptor T cells is disorganized and does not form a bull’s eye structure.^5^ Multiple Lck microclusters have been observed with chimeric antigen receptor T cells, in contrast to the typical Lck clustering at the central supramolecular activation cluster region.^5^

The density of antigen epitopes is critical for chimeric antigen receptor T activation. A threshold of antigen density exists such that chimeric antigen receptor T cells are activated only after it reaches a threshold amount. Stone et al. proposed a “sensitivity scale” model for targeting cells with chimeric antigen receptors and bispecific T cell engagers.^6^ The threshold required for cytokine secretion is significantly higher than that for cytolytic activity. The molecular properties of chimeric antigen receptor T cell targets can influence chimeric antigen receptor signaling. For example, the more proximal the target epitope of CD20 is to the membrane, the longer the length of the chimeric antigen receptor’s hinge/spacer required to optimize the receptor-ligand binding.^7-10^ Conversely, a longer target epitope achieves optimal stimulation of chimeric antigen receptors by requiring a shorter length of the chimeric antigen receptor’s hinge/spacer.

The first successful target for chimeric antigen receptor T therapy, CD19, is a glycosylphosphatidylinositol-anchored protein. The glycosylphosphatidylinositol anchor is a posttranslational modification of protein with a glycolipid. The carboxyl terminus of all glycosylphosphatidylinositol-anchored proteins contains a hydrophobic signal sequence that triggers the addition of the glycosylphosphatidylinositol anchor. Glycosylphosphatidylinositol-anchored proteins are transported to the cell surface after the carboxyl-terminal stretch of hydrophobic amino acids is clipped off and replaced with a preassembled glycosylphosphatidylinositol anchor. Glycosylphosphatidylinositol-anchored proteins may transduce signals for cell proliferation and motility, and they are known to be associated with lipid rafts enriched in sphingolipids and cholesterol.

To investigate whether the arrangement of epitopes on an antigen affects chimeric antigen receptor T cell activation, we grafted a well-characterized MUC1 epitope^11-14^ (recognized by 16A-chimeric antigen receptor) to the glycosylphosphatidylinositol anchor protein CD24. We then compared the stimulation of chimeric antigen receptor T cells by a single target epitope displayed as parallel pillars versus consecutive tandem repeating epitopes and found increased stimulation with the parallel arrangement.

## Methods

### Stable expression of different MUC1 molecules in 293T-COSMC^-/-^ cells

To construct plasmid pMUC1(1TR), we synthesized complementary DNA sequences encoding the signal peptide, the variable number of tandem repeat (one tandem repeat [1TR]) region, transmembrane region, and cytoplasmic domain of the MUC1 molecule and cloned them into a pcDNA3.1 plasmid (Thermo Fisher, Waltham, MA).

To construct plasmid pCD24-MUC1(1TR), we synthesized complementary DNA sequences encoding the signal peptide of CD24 and the variable number of tandem repeat (1TR) region fused to a peptide core of the CD24 molecule and cloned them into a pcDNA3.1 plasmid.

To construct plasmid pCD24-MUC1(5TR), we synthesized complementary DNA sequences encoding the signal peptide of CD24 and the variable number of tandem repeat (5TR) region fused to a peptide core of the CD24 molecule and cloned them into a pcDNA3.1 plasmid. The sequences of our plasmids are listed in Supplementary Figure 1.

Different MUC1 molecules were stably transfected into 293T-COSMC^-/-^ cells (published elsewhere, provided by Prof. Yan Zhang, Shanghai Jiaotong University) by Lipofectamine (Thermo Fischer) transfection and selection with hygromycin-containing culture medium. The MUC1-positive cells were obtained by flow cytometry–based cell sorting, with similar mean fluorescence intensity measured by 16A antibody staining (BioLegend, San Diego, CA). The mean fluorescence intensity for MUC1(1TR), CD24-MUC1(1TR), and CD24-MUC1(5TR) were 1.08E6, 4.98E5, and 7.74E5, respectively.

### Preparation of recombinant lentivirus encoding chimeric antigen receptors

The pHAGE-16A-chimeric antigen receptor-red fluorescent protein (RFP) plasmid was constructed by assembling complementary DNA sequences of the heavy chain variable domain (V_H_), hinge region (GGGGSGGGGSGGGGS), light chain variable domain (V_L_), transmembrane region of CD8α, intracellular domain of 41BB, and CD3. The sequences of the plasmids are listed in Supplementary Figure 2.

To generate viral particles, Lenti-293X cells (Sinobiological, China) were transfected with pHAGE 16A-chimeric antigen receptor-RFP plasmid or mock pHAGE-IRES-RFP plasmid, together with packaging plasmids psPAX2 and pMD2.G. Then viral particles were collected and concentrated 100-fold by mixing with PEG 8000 (Sigma-Aldrich, St Louis, MO) for 12 h at 4°C and centrifuging for 30 min at 1600×g at 4°C.

### Generation of chimeric antigen receptor T cells

Peripheral blood mononuclear cells from healthy donors were activated in cell culture plates precoated with CD3 (5 μg/mL, BioLegend, San Diego, CA) and CD28 (1 μg/mL, BD Biosciences, San Jose, CA) antibodies in complete human lymphocyte culture medium (Gibco RPMI Medium 1640), 10% Gibco heat-inactivated fetal bovine serum, 100 units/mL penicillin, and 100 μg/mL streptomycin, with 200 IU/mL recombinant human interleukin-2 (SI HUAN SHENG WU, Beijing, China). After 48 h, peripheral blood mononuclear cells were infected with viral supernatant at a multiplicity of infection of 10 to 40. Empty vectors of pHAGE were also cotransduced into T cells to generate control T cells. After 12 h, cells were fed with fresh completed medium and then expanded within 10 days before functional experiments.

### Stimulation of chimeric antigen receptor T cells

The chimeric antigen receptor T cells were mixed with 100,000 MUC1-expressing target cells at the effector/target ratio of 1:2. Chimeric antigen receptor T cells and MUC1-expressing target cells were co-cultured in 96-well plates for 24 hours, and the cytokines released to culture medium were measured by enzyme-linked immunosorbent assay. As a positive control, phorbol myristate acetate (500 ng/mL) and ionomycin (10 μg/mL, both from Dakewe Biotech Co., China) were used to stimulate chimeric antigen receptor T cells.

## Results

### MUC1 epitope presented on glycosylphosphatidylinositol anchor showed greater stimulation of chimeric antigen receptor T cells

To test our hypothesis that a glycosylphosphatidylinositol anchor–carried epitope may significantly influence the stimulation of chimeric antigen receptor T cells, we designed the CD24-MUC1-TR antigen, in which a tandem repeat sequence of MUC1 is fused with CD24 protein, and compared its stimulatory activity to a MUC1 molecule with the original transmembrane domain (MUC1-TR). As measured by enzyme-linked immunosorbent assay, a 6-to 20-fold increase was observed when CD24-MUC1-TR cells were used to stimulate cytokine production by chimeric antigen receptor T cells, compared with the MUC1-TR molecule, which uses the original transmembrane domain of MUC1.

### MUC1 epitope presented as parallel lipid-carried epitope pillars increased stimulation of chimeric antigen receptor T cells

To compare the stimulatory activity of epitopes displayed as parallel lipid-carried epitope pillars versus lipid-carried tandem repeat sequences, we designed CD24-5MUC1 molecule, which contains five consecutive tandem repeat sequences, and compared its stimulatory activity with CD24-MUC1-TR, which contains only one epitope displayed on the cell surface as pillars in parallel. As measured by enzyme-linked immunosorbent assay, a greater than 10-fold decrease in stimulation by CD24-5MUC1 was observed as compared with CD24-MUC1-TR.

## Discussion

### Abnormally glycosylated glycosylphosphatidylinositol anchor protein as target for chimeric antigen receptor T therapy

Our results indicated that lipid-carried epitope pillars in parallel are ideal targets for chimeric antigen receptor T therapy. Glycosylphosphatidylinositol-anchored proteins, such as CD19 and CD20, have been targets for several chimeric antigen receptor T therapies approved by the US Food and Drug Administration. However, healthy B cells expressing CD19 or CD20 are also eliminated by current chimeric antigen receptor T cells used clinically. Abnormal O-glycosylation has been reported in multiple types of cancer, including leukemia and lymphoma. Chimeric antigen receptor T cells have been designed to distinguish truncated and normal glycoforms, with increased antitumor efficacy but decreased toxicity to nontumoral tissue.^15^ Neurotoxicity is a severe, even fatal, adverse event due to nonspecific binding to normal CD19 expressed by brain mural cells.^16^ However, such toxicity would be avoided with more specific chimeric antigen receptors targeting abnormally glycosylated CD19. We and others have developed monoclonal antibodies and chimeric antigen receptor T cells targeting abnormally glycosylated MUC1 glycopeptides.^11^ Antibodies targeting a glycopeptide epitope of abnormally glycosylated glycosylphosphatidylinositol-anchored proteins may be developed by similar strategies and bring about a new panoply of tumor-specific antibodies and chimeric antigen receptors.

### Epitope pillars in parallel are superior to consecutively aligned epitope tandem repeats in stimulating chimeric antigen receptor T cells

Both the epitopes displayed on CD19 and CD20 molecules are single epitopes, whereas many other tumor antigens such as MUC1 contain repeating units of epitopes in a single molecule. The loss of stimulation of chimeric antigen receptor T cells by target cells expressing MUC1 tandem repeats indicates that such consecutively aligned epitopes may cause inefficient binding of chimeric antigen receptors to the cell surface. The exact mechanisms for such loss of stimulation remain to be studied, but here we suggest some possible scenarios (Figure 3). Firstly, the clustering of chimeric antigen receptors around the tandem repeat area might be inefficient because the tandem repeat region of mucin typically is considered as disordered and lacks alpha-helix or beta-sheet secondary structures. Secondly, the mechanosensing of epitope pillars in parallel may generate increased signaling strength than consecutively aligned epitopes because lipid-carried epitopes displayed in parallel are more elastic than those displayed as tandem repeats. Thirdly, the protein segregation of the immune synapse around the epitope pillars in parallel may be faster than consecutively aligned epitopes. All of these proposed mechanisms are worthy of further studies at the single molecule level and may provide clues for better selection of targets for chimeric antigen receptor T stimulation. Imaging methods have been developed to measure synapse quality that may predict clinical outcome of chimeric antigen receptor T therapy; these include quantitation of clustering of tumor antigen and distribution of key signaling molecules within the immunological synapse.^17^ Artificially designed antigen molecules with well-characterized tumor epitope structures may be designed as calibrators to measure the speed of epitope movement and the elasticity that determines force-regulated mechanosensing of chimeric antigen receptors.

**Figure 1.**
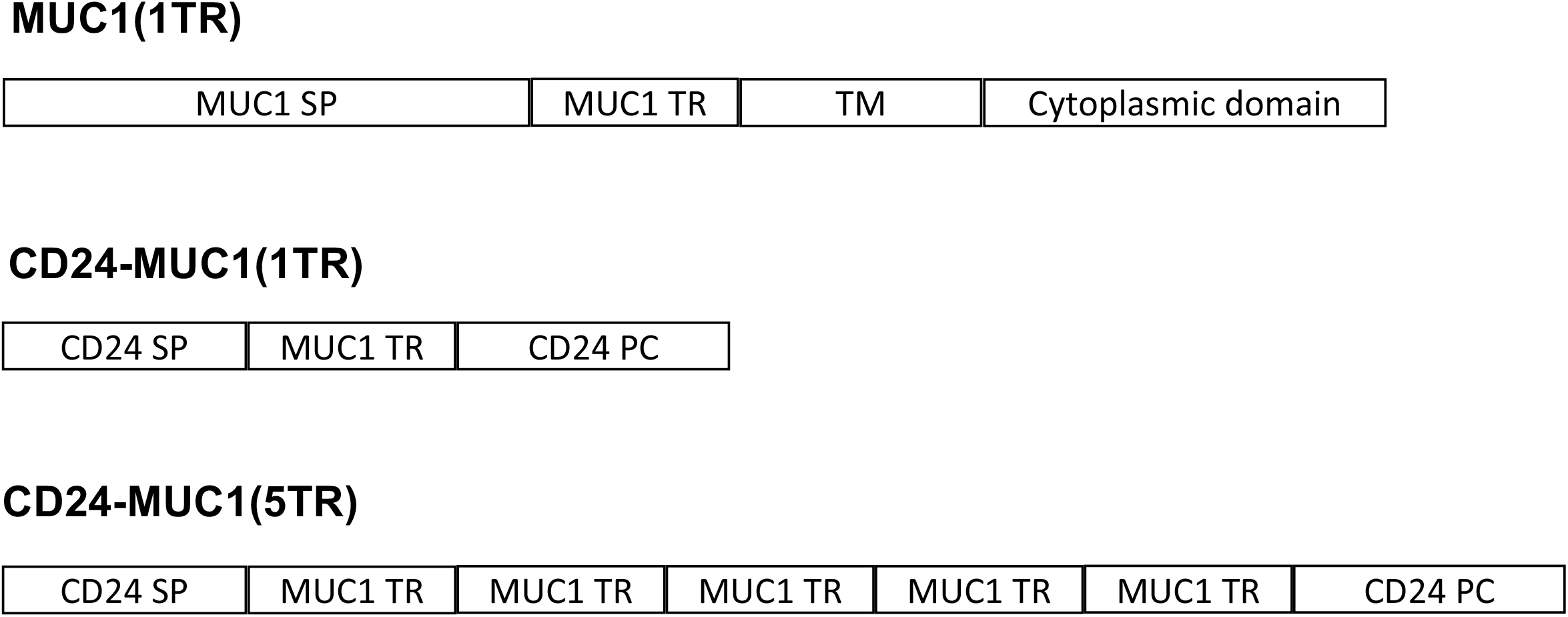
Structure of MUC1 antigens in this study. MUC1(1TR) contains the original signal peptide (MUC1 SP) of MUC1 molecule, one tandem repeat sequence of MUC1 (MUC1 TR), a transmembrane domain (TM), and the cytoplasmic domain of the original MUC1 molecule. CD24-MUC1(1TR) contains the signal peptide of CD24 (CD24 SP), one tandem repeat sequence of MUC1 (MUC1 TR), and the protein core of the CD24 molecule (CD24 PC). CD24-MUC1(5TR) contains the signal peptide of CD24 (CD24 SP), five tandem repeat sequences of MUC1 (MUC1 TR), and the protein core of the CD24 molecule (CD24 PC).

**Figure 2.**
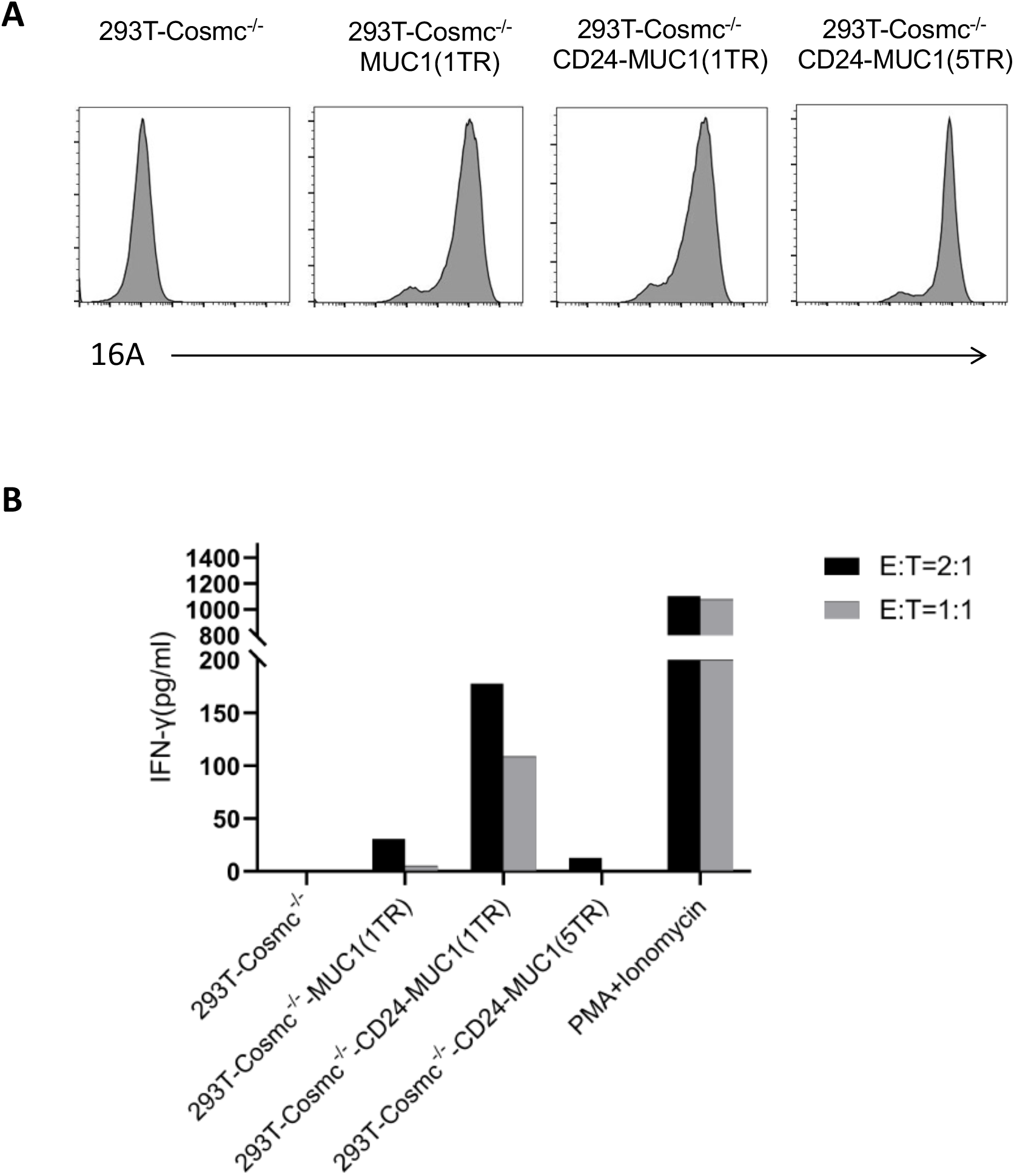
Stimulation of 16A-chimeric antigen receptor T cells by different MUC1 molecules. A. Different MUC1 molecules were stably expressed in 293T-COMSC^-/-^ cells. The MUC1-positive cells were obtained by flow cytometry–based cell sorting, with similar mean fluorescence intensity measured by 16A antibody staining. The mean fluorescence intensity for MUC1(1TR), CD24-MUC1(1TR), and CD24-MUC1(5TR) were 1.08E6, 4.98E5, and 7.74E5, respectively. (See Fig. 1 for descriptions of the MUC1 samples’ names and structures.) B. 16A-chimeric antigen receptor T cells were stimulated by 293T-COSMC^-/-^ cells expressing different MUC1 molecules using an effector (E)/target (T) ratio of 2:1, respectively. The secretion of interferon (IFN)-γ by the chimeric antigen receptor T cells was measured by enzyme-linked immunosorbent assay.

**Figure 3.**
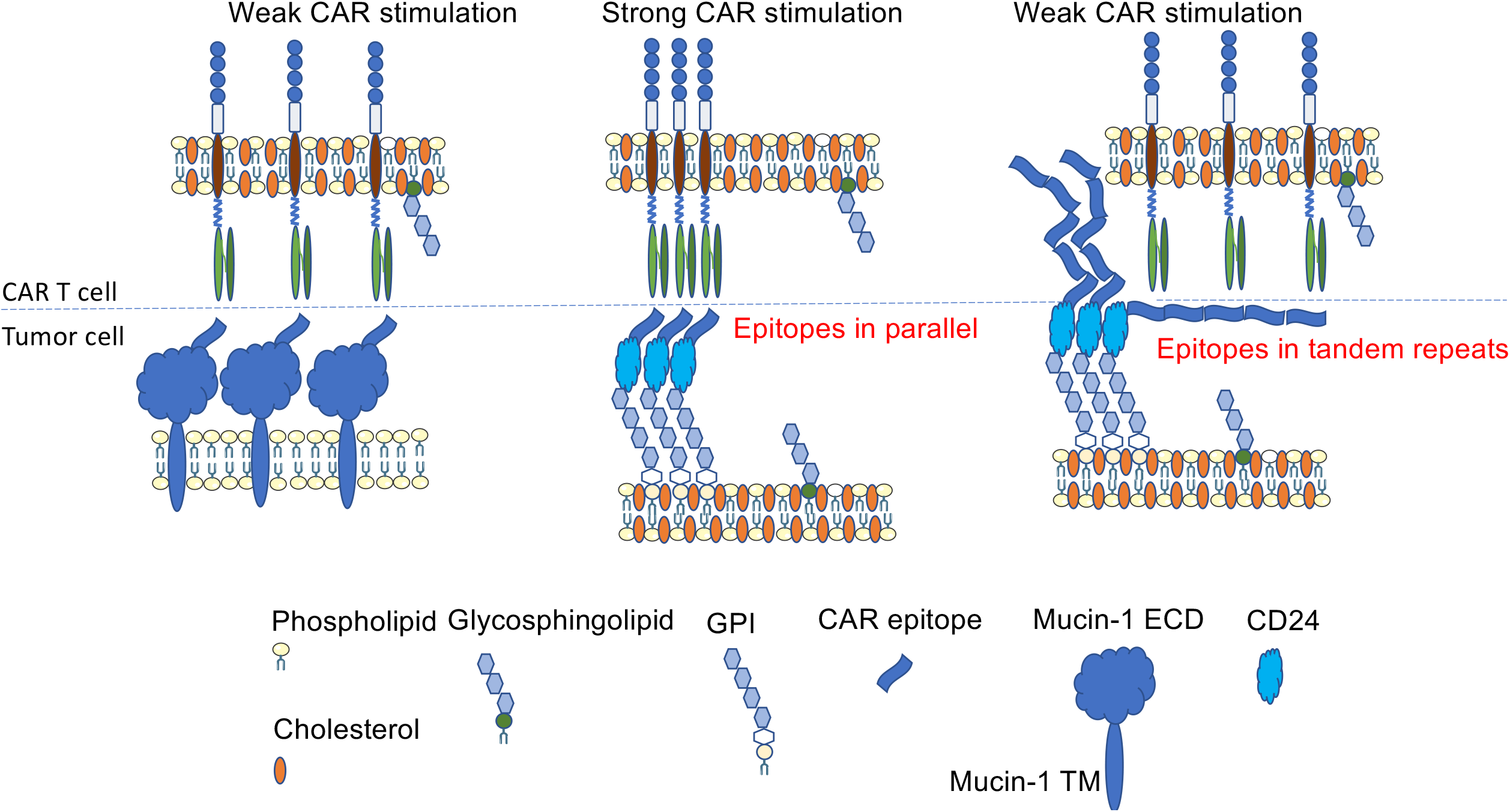
Parallel lipid-carried epitope model for chimeric antigen receptor T therapy. As shown in the central panel, strong stimulation to chimeric antigen receptor T cells is achieved when tumor antigen epitopes are carried by lipids located on lipid rafts containing cholesterol and glycosphingolipids. When tumor antigen epitopes are aligned as tandem repeats (right panel), stimulation is drastically reduced. When tumor antigen epitopes are carried by glycoproteins (left panel), stimulation is less efficient as compared with the same epitopes carried by lipids.

Past studies on the epitopes of chimeric antigen receptor T therapy have found that the density of epitopes is critical for chimeric antigen receptor T stimulation. For example, a “sensitivity scale” was proposed for a tumor-specific glycopeptide epitope on the sialomucin-like transmembrane glycoprotein OTS8.^6^ Our studies clearly showed that the carriers of epitopes are critical, and lipid carriers facilitate the assembly of the epitope patch required for chimeric antigen receptor T stimulation. Past studies also showed that the closer arrangement of tumor antigen epitope to membrane required a longer version of the chimeric antigen receptor’s hinge region.^8^ In our study, we have used a longer version of hinge region for the chimeric antigen receptor, but it did not improve the weak stimulation of the 16A epitope arranged as tandem repeats on lipid carriers (Figure 3, right panel) or arranged in parallel on protein carriers (Figure 3, left panel).

Currently, there are several limitations to explore the mechanisms of parallel lipid-carried epitopes. To measure the mechanosensing of chimeric antigen receptors, parallel lipid-carried epitopes should be synthesized as tumor antigen proteins inserted on lipid rafts. To measure the speed of epitope movement in lipid rafts, proper single molecule imaging instrument and reagents are required. There have also been reports that glycoproteins such as MUC1 may be distributed in lipid rafts.^18^ The percentage of MUC1 distributed in lipid rafts should be determined by mass spectrometry, when studying the interactions of chimeric antigen receptor T cells with MUC1 located in lipid rafts versus those outside of lipid rafts. Similarly, a tumor epitope may be expressed as both lipid-carried and protein-carried, such as the sialic acid epitope on ganglioside GD2, and on glycoprotein CD166.^19^ The percentage of epitopes carried by gangliosides should be determined and compared with those carried by CD166, when our parallel lipid-carried model is applied to the tumor cells.

Our findings and the parallel lipid-carried epitope model provide a clearer and fine-tuned view for the epitope arrangement of tumor antigen, that will precisely discriminate the tumor cells from normal healthy cells. Those epitopes fitting the parallel lipid-carried epitope model in tumor cells will be a major type of functional epitopes to stimulate chimeric antigen receptor T cells. To avoid deadly cytokine storm and toxicity to normal cells, parallel lipid-carried epitopes should be highly tumor-specific, such as those targeting abnormal glycosylation of tumor cells.

## Author contributions

JT, HL, and DZ designed this study. JT, HL, JW, CZ, XM and DZ contributed to the collection, analysis, and interpretation of data. DZ wrote the manuscript. All authors read and approved the final manuscript.

## Funding

This work was supported by National Key Research and Development Plan grants 2021YEE0200500 and 2017YFA0505901, Fundamental Research Funds for the Central Universities 22120200163, National Natural Science Foundation of China grant 31870972, and Shanghai Science and Technology Commission grant 15002360172.

All these sponsors have no roles in the study design or the collection, analysis, and interpretation of data.

### Abbreviations

CAR: Chimeric antigen receptor
COSMC: (C1GALT1C1)
C1GALT1: Specific Chaperone 1
GPI: Glycosylphosphatidylinositol
GSL: glycosphingolipid
MUC1: Mucin-1
TR: tandem repeat

---

**Figure S1.**
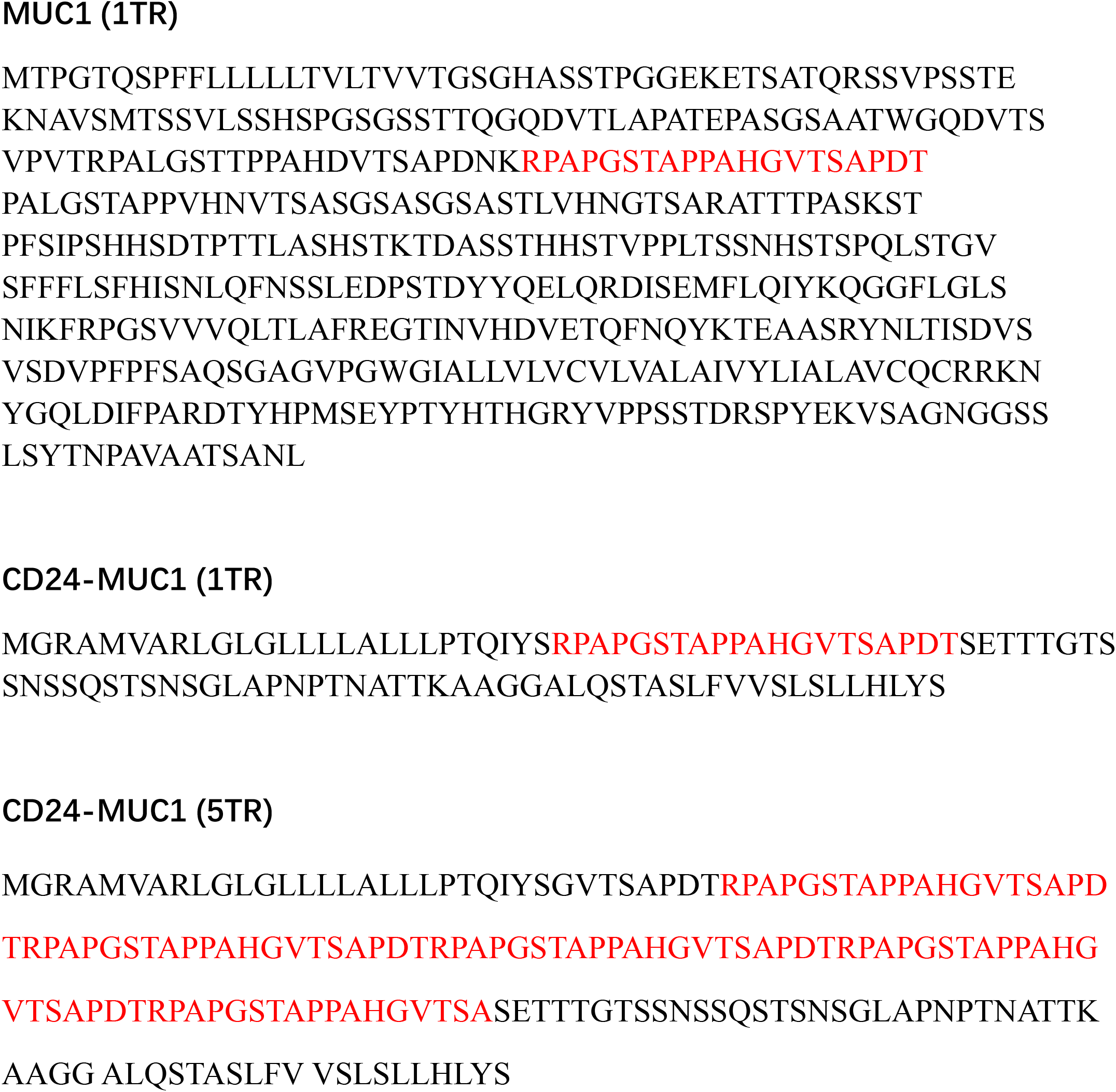
Amino acid sequences of different MUC1 molecules.

**Figure S2.**
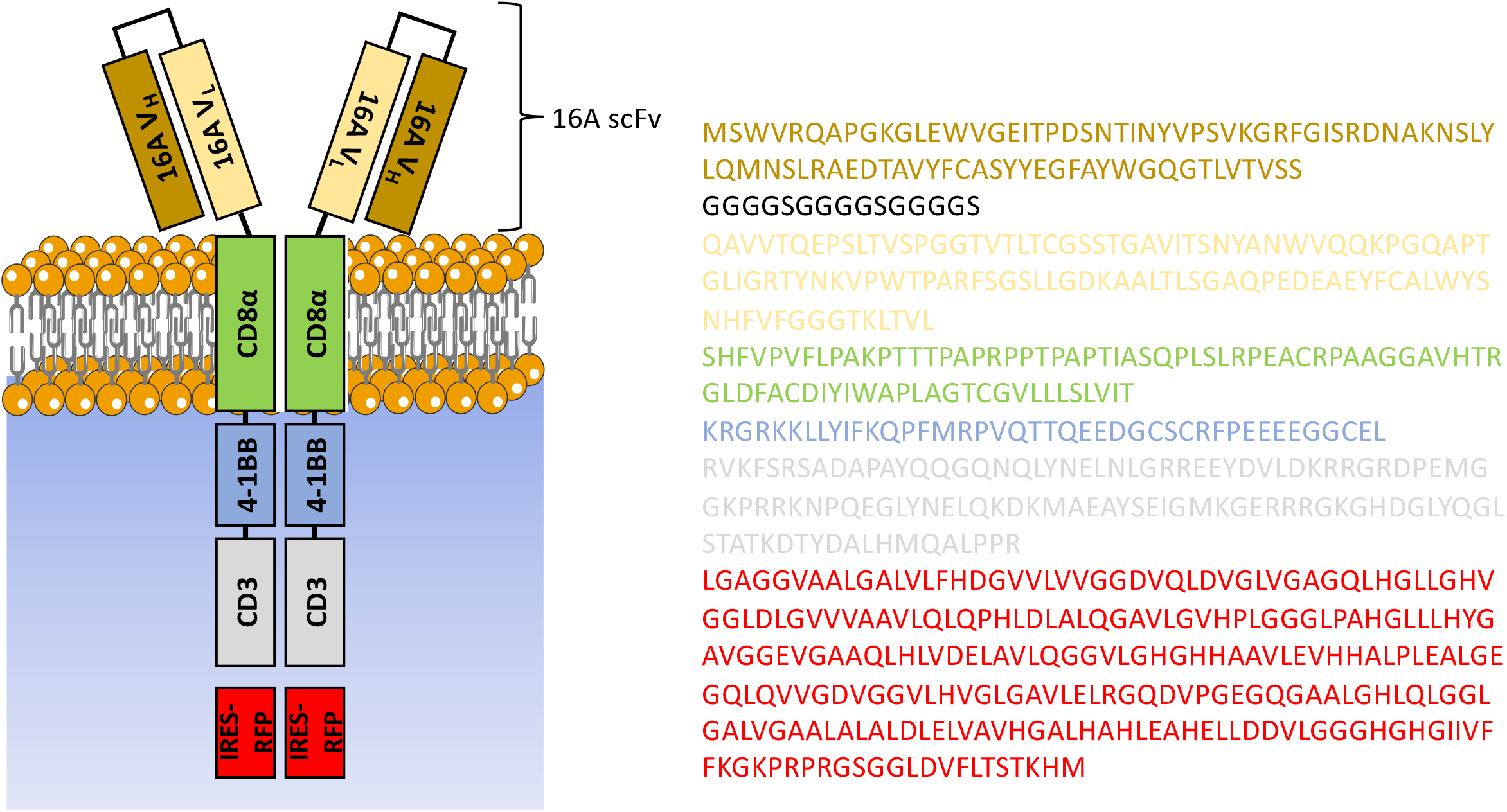
Amino acid sequence of 16A chimeric antigen receptor.

## Notes

**Conflict of interest disclosures** The authors have no conflict of interest to declare

### Competing Interest Statement

The authors have declared no competing interest.

## References

1. Wu L, Wei Q, Brzostek J, Gascoigne NRJ. Signaling from T cell receptors (TCRs) and chimeric antigen receptors (CARs) on T cells. Cell Mol Immunol. 2020 Jun;17(6):600–612.

2. Salter AI, Ivey RG, Kennedy JJ, Voillet V, Rajan A, Alderman EJ, Voytovich UJ, Lin C, Sommermeyer D, Liu L, Whiteaker JR, Gottardo R, Paulovich AG, Riddell SR. Phosphoproteomic analysis of chimeric antigen receptor signaling reveals kinetic and quantitative differences that affect cell function. Sci Signal. 2018 Aug 21;11(544):eaat6753.

3. Harris DT, Hager MV, Smith SN, Cai Q, Stone JD, Kruger P, Lever M, Dushek O, Schmitt TM, Greenberg PD, Kranz DM. Comparison of T Cell Activities Mediated by Human TCRs and CARs That Use the Same Recognition Domains. J Immunol. 2018 Feb 1;200(3):1088–1100.

4. Watanabe K, Kuramitsu, Posey AD, and June CH. Expanding the Therapeutic Window for CAR T Cell Therapy in Solid Tumors: The Knowns and Unknowns of CAR T Cell Biology. Front Immunol. 2018 Oct 26;9:2486.

5. Xiong W, Chen Y, Kang X, Chen Z, Zheng P, Hsu YH, Jang JH, Qin L, Liu H, Dotti G, Liu D. Immunological synapse and CAR Immunological Synapse Predicts Effectiveness of Chimeric Antigen Receptor Cells. Mol Ther. 2018 Apr 4;26(4):963–975.

6. Stone JD, Aggen DH, Schietinger A, Schreiber H, Kranz DM. A sensitivity scale for targeting T cells with chimeric antigen receptors (CARs) and bispecific T-cell Engagers (BiTEs). Oncoimmunology. 2012 Sep 1;1(6):863–873.

7. Hombach AA, Schildgen V, Heuser C, Finnern R, Gilham DE, Abken H. T cell activation by antibody-like immunoreceptors: the position of the binding epitope within the target molecule determines the efficiency of activation of redirected T cells. J Immunol. 2007 Apr 1;178(7):4650–7.

8. James SE, Greenberg PD, Jensen MC, Lin Y, Wang J, Till BG, Raubitschek AA, Forman SJ, Press OW. Antigen sensitivity of CD22-specific chimeric TCR is modulated by target epitope distance from the cell membrane. J Immunol. 2008 May 15;180(10):7028–38.

9. Zah E, Lin MY, Silva-Benedict A, Jensen MC, Chen YY. T Cells Expressing CD19/CD20 Bispecific Chimeric Antigen Receptors Prevent Antigen Escape by Malignant B Cells. Cancer Immunol Res. 2016 Jun;4(6):498–508.

10. Qin H, Cho M, Haso W, Zhang L, Tasian SK, Oo HZ, Negri GL, Lin Y, Zou J, Mallon BS, Maude S, Teachey DT, Barrett DM, Orentas RJ, Daugaard M, Sorensen PH, Grupp SA, Fry TJ. Eradication of B-ALL using chimeric antigen receptor-expressing T cells targeting the TSLPR oncoprotein. Blood. 2015 Jul 30;126(5):629–39.

11. Zhou D, Xu L, Huang W, Tonn T. Molecules Epitopes of MUC1 Tandem Repeats in Cancer as Revealed by Antibody Crystallography: Toward Glycopeptide Signature-Guided Therapy. Molecules. 2018 May 31;23(6):1326.

12. Song W, Delyria ES, Chen J, Huang W, Lee JS, Mittendorf EA, Ibrahim N, Radvanyi LG, Li Y, Lu H, Xu H, Shi Y, Wang LX, Ross JA, Rodrigues SP, Almeida IC, Yang X, Qu J, Schocker NS, Michael K, Zhou D. MUC1 glycopeptide epitopes predicted by computational glycomics. Int J Oncol. 2012 Dec;41(6):1977–84.

13. Zhou D, Xiao K, Tian Z. Separation and detection of minimal length glycopeptide neoantigen epitopes centering the GSTA region of MUC1 by liquid chromatography/mass spectrometry. Rapid Commun Mass Spectrom. 2020 Mar 30;34(6):e8622.

14. Pan D, Tang Y, Tong J, Xie C, Chen J, Feng C, Hwu P, Huang W, Zhou D. Cancer Medicine An antibody-drug conjugate targeting a GSTA glycosite-signature epitope of MUC1 expressed by non-small cell lung cancer. Cancer Med. 2020 Dec;9(24):9529–9540.

15. Posey AD Jr, Schwab RD, Boesteanu AC, Steentoft C, Mandel U, Engels B, Stone JD, Madsen TD, Schreiber K, Haines KM, Cogdill AP, Chen TJ, Song D, Scholler J, Kranz DM, Feldman MD, Young R, Keith B, Schreiber H, Clausen H, Johnson LA, June CH. Engineered CAR T Cells Targeting the Cancer-Associated Tn-Glycoform of the Membrane Mucin MUC1 Control Adenocarcinoma. Immunity. 2016 Jun 21;44(6):1444–54.

16. Kevin R. Parker, Denis Migliorini, Eric Perkey, Kathryn E. Yost, Aparna Bhaduri, Puneet Bagga, Mohammad Haris, Neil E. Wilson, Fang Liu, Khatuna Gabunia, John Scholler, Thomas J. Montine, Vijay G. Bhoj, Ravinder Reddy, Suyash Mohan, Ivan Maillard, Arnold R. Kriegstein, Carl H. June, Howard Y. Chang, Avery D. Posey, Ansuman T. Satpathy. Single-Cell Analyses Identify Brain Mural Cells Expressing CD19 as Potential Off-Tumor Targets for CAR-T Immunotherapies, Cell. 2020. Oct 1; 183(1):126–142.

17. Xiong W, Chen Y, Kang X, Chen Z, Zheng P, Hsu YH, Jang JH, Qin L, Liu H, Dotti G, Liu D. Immunological Synapse Predicts Effectiveness of Chimeric Antigen Receptor Cells. Mol Ther. 2018 Apr 4;26(4):963–975.

18. Staubach S, Razawi H, Hanisch FG. Proteomics of MUC1-containing lipid rafts from plasma membranes and exosomes of human breast carcinoma cells MCF-7. Proteomics. 2009 May;9(10):2820–35.

19. Agrawal, V., & Frankel, A. E. (2010). 14G2a anti-GD2 crossreactivity with the CD166 antigen. Journal of Immunotherapy, 33(9), 1014–1015.

